# High-Frequency Stimulation Does Not Improve Comfort of Transcutaneous Spinal Cord Stimulation

**DOI:** 10.1101/2022.09.06.506783

**Authors:** Ashley N Dalrymple, Charli Ann Hooper, Minna G Kuriakose, Marco Capogrosso, Douglas J Weber

**Affiliations:** Department of Mechanical Engineering, Carnegie Mellon University, Pittsburgh, PA, USA; NeuroMechantronics Lab, Carnegie Mellon University, Pittsburgh, PA, USA; Department of Biomedical Engineering, Carnegie Mellon University, Pittsburgh, PA, USA; Department of Bioengineering, University of Pittsburgh, Pittsburgh, PA, USA; Department of Neurological Surgery, University of Pittsburgh, Pittsburgh, PA, USA; Rehab Neural Engineering Labs, University of Pittsburgh, Pittsburgh, PA, USA; Center for Neural Basis of Cognition, Pittsburgh, PA, USA; Neuroscience Institute, Carnegie Mellon University, Pittsburgh, PA, USA

**Keywords:** Transcutaneous spinal cord stimulation, high-frequency stimulation, neuromodulation, reflexes, pain tolerance

## Abstract

Spinal cord neuromodulation has gained much attention for demonstrating improved motor recovery in people with spinal cord injury, motivating the development of clinically applicable technologies. Among them, transcutaneous spinal cord stimulation (tSCS) is attractive because of its non-invasive profile. Many tSCS studies employ a high-frequency (10 kHz) carrier, which has been reported to reduce stimulation discomfort. However, these claims have come under scrutiny in recent years. The purpose of this study was to determine whether high-frequency tSCS is more comfortable at therapeutic amplitudes, which evoke posterior root-muscle (PRM) reflexes. In 16 neurologically intact participants, tSCS was delivered using a 1-ms long monophasic pulse with and without a high-frequency carrier. Stimulation amplitude and pulse duration were varied and PRM reflexes were recorded from the soleus, gastrocnemius, and tibialis anterior muscles. Participants rated their discomfort during stimulation from 0-10 at PRM reflex threshold. At PRM reflex threshold, high-frequency stimulation (0.87 ± 0.2) was equally comfortable as conventional stimulation (1.03 ± 0.18) but required approximately double the charge to evoke the PRM reflex (conventional: 32.4 ± 9.2 µC; high-frequency: 62.5 ± 11.1 µC). Strength-duration curves for high-frequency stimulation had a rheobase that was 4.8X greater and a chronaxie that was 5.7X narrower than the conventional monophasic pulse, indicating that high-frequency stimulation was less efficient in recruiting neural activity in spinal roots. High-frequency tSCS is equally as comfortable as conventional stimulation at amplitudes required to stimulate spinal dorsal roots.

**GRAPHICAL ABSTRACT:** 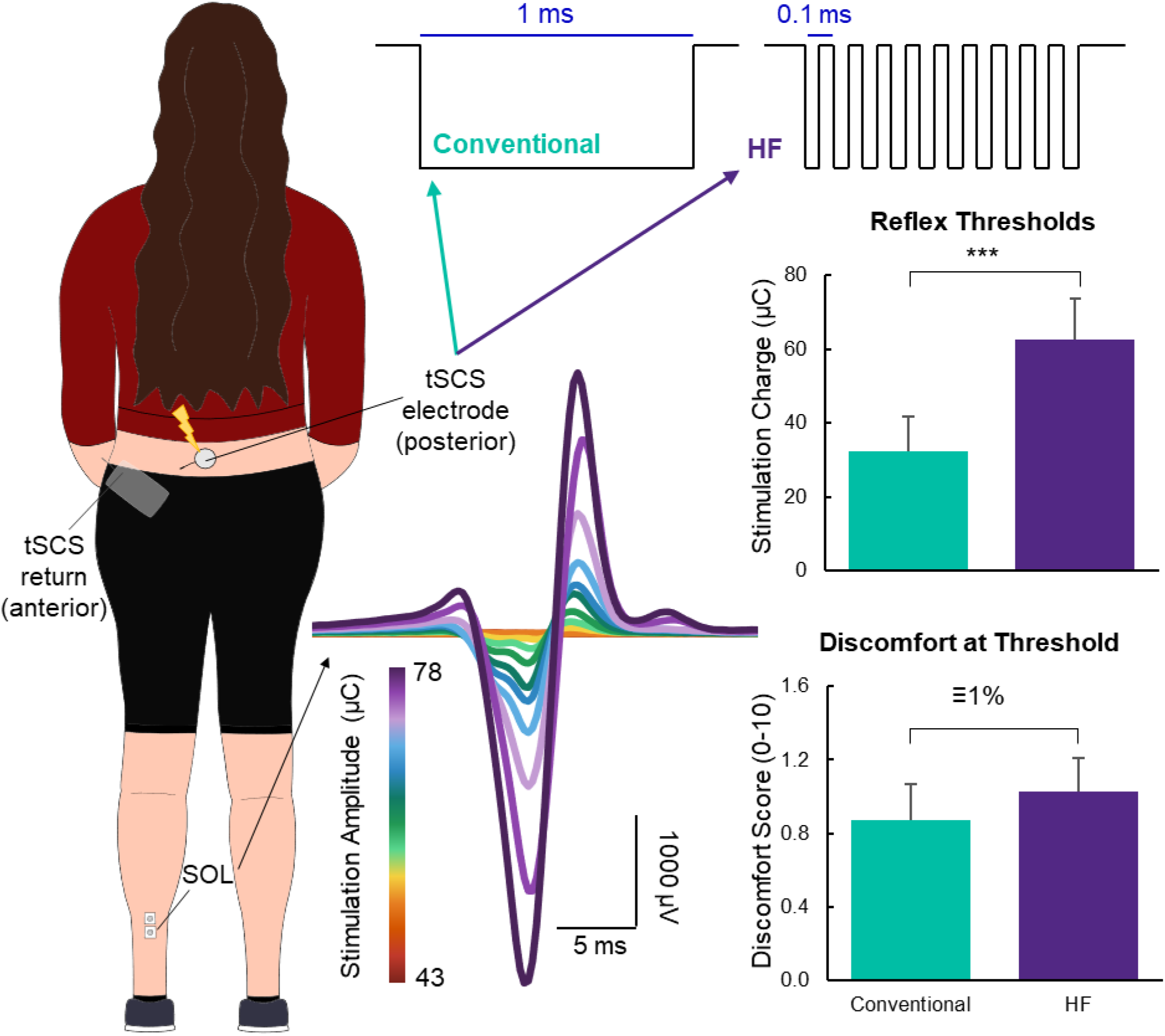

## INTRODUCTION

Neuromodulation of the spinal cord can be used to treat chronic pain (Caylor et al., 2019; Shealy et al., 1967) and improve motor function after stroke and spinal cord injury (Angeli et al., 2018; Carhart et al., 2004; Gill et al., 2018; Harkema et al., 2011; Powell et al., 2022; Rowald et al., 2022). Transcutaneous spinal cord stimulation (tSCS) is a non-invasive neuromodulation technique that uses adhesive electrodes on or adjacent to the spine to activate primary afferent neurons in the spinal roots, particularly the large diameter afferent fibers, similar to SCS (Capogrosso et al., 2013; Hofstoetter et al., 2018). tSCS is a more accessible neuromodulation therapy that could potentially reach people without access to surgical procedures or those who are contra-indicated or unwilling to receive a SCS implant. To date, tSCS has been demonstrated as an effective method to improve motor function in the arm and hand (Freyvert et al., 2018; Gad et al., 2018; Inanici et al., 2021, 2018), trunk (Keller et al., 2021; Rath et al., 2018; Sayenko et al., 2019), and legs (Gad et al., 2017, 2019; Hofstoetter et al., 2013, 2015b, 2021; Samejima et al., 2022) in people with various neurological conditions.

tSCS (and SCS) excite the large-diameter afferents in the spinal roots (Hofstoetter et al., 2019, 2018), engaging spinal reflex pathways that facilitate activity in motoneurons. The evoked response can be recorded using electromyography (EMG) and is known as the posterior root-muscle (PRM) reflex (Minassian et al., 2007). The PRM reflex is equivalent to the Hoffman (H)- reflex but engages multiple segments of spinal roots (Krenn et al., 2013; Minassian et al., 2007). Compared to SCS with implanted electrodes, the current amplitudes required for tSCS are much higher. Since tSCS is delivered through electrodes placed on the skin, stimulation also activates cutaneous afferents and induces strong contractions in paraspinal muscles, both of which can cause discomfort.

The conventional waveform used for tSCS is a square pulse with a duration of 1-2 ms applied at 30-50 Hz to facilitate muscle activation (Freyvert et al., 2018; Gad et al., 2017; Hofstoetter et al., 2014, 2015b, 2020, 2021; Knikou and Murray, 2019). An alternative approach uses a high frequency carrier wave, typically 10 kHz, within the 1-2 ms pulse (Gad et al., 2019, 2018; Gerasimenko et al., 2016; Y. Gerasimenko et al., 2015; Y. P. Gerasimenko et al., 2015; Inanici et al., 2021, 2018; Keller et al., 2021; Rath et al., 2018; Sayenko et al., 2019). The concept of using a high-frequency carrier originates from Russia in the 1970’s and is colloquially termed “Russian current” (Manson et al., 2020; Ward and Shkuratova, 2002). The benefit of the high-frequency carrier, originally selected to be 2.5 kHz, was that it made the stimulation less painful. A later study found the optimal frequency for motor activation and minimal pain to be 10 kHz (Ward and Robertson, 1998). Several tSCS studies claim that stimulation at 10 kHz is painless due to blocking of superficial nociceptive afferents, enabling use of higher stimulation amplitudes, such as those needed to target the spinal roots (Gad et al., 2017; Y. Gerasimenko et al., 2015; Inanici et al., 2021; Manson et al., 2020; Sayenko et al., 2019).

A recent study aimed to characterize the maximal tolerable stimulation intensity when using a conventional biphasic stimulation waveform with and without a 5 kHz carrier frequency (Manson et al., 2020). They found that when the maximum tolerable stimulation intensity was normalized to the stimulation intensity required to evoke a PRM reflex, there was no difference between the waveforms. It is important to note that tSCS for neuromodulation to facilitate motor functions requires stimulation amplitudes near the PRM reflex threshold to be effective. Therefore, characterizing the comfort of tSCS at the PRM reflex threshold is of paramount interest.

Here, we compared stimulation thresholds for evoking PRM reflexes using a conventional 1 ms-long monophasic pulse with and without a 10 kHz carrier frequency for tSCS. Participants were asked to report a discomfort rating for the stimulation applied at the reflex threshold. We hypothesized that high-frequency stimulation would have a higher PRM reflex threshold, and that at threshold, the high-frequency waveform would not be more comfortable than the conventional waveform. As hypothesized, PRM reflex thresholds were higher for the high-frequency waveform, which was equally as comfortable as the conventional waveform at its PRM reflex threshold. These results suggest that for tSCS, a high-frequency waveform offers no advantages for reducing discomfort compared to a conventional waveform. Moreover, the higher stimulation charge needed to evoke a response with the high-frequency waveform requires the stimulator to produce much higher voltages, increasing the power budget and risk of injury.

## MATERIALS AND METHODS

### Participants

Sixteen neurologically intact individuals participated in this study (mean age ± standard deviation (SD): 29.5 ± 6.9 years; 8 female; Table 1). We measured the circumference of each participant’s waist and hips.

**Table 1.** Demographic information for research participants. All participants were neurologically intact. HF = high-frequency.

| Participant ID | Sex | Age (years) | Height (cm) | Weight (lbs) | Waist (cm) | Hip (cm) | First Waveform |
| --- | --- | --- | --- | --- | --- | --- | --- |
| TSP01 | Male | 49 | 180 | 195 | 102 | 104 | Conventional |
| TSP02 | Female | 23 | 165 | 125 | 69 | 92 | HF |
| TSP03 | Female | 25 | 160 | 119 | 71.5 | 90 | Conventional |
| TSP04 | Female | 26 | 160 | 108 | 69 | 80 | Conventional |
| TSP05 | Male | 28 | 193 | 190 | 91 | 97 | HF |
| TSP06 | Male | 40 | 178 | 170 | 86.5 | 93 | Conventional |
| TSP07 | Male | 24 | 180 | 139 | 76 | 92 | HF |
| TSP08 | Female | 23 | 168 | 135 | 69.5 | 88.5 | Conventional |
| TSP09 | Male | 24 | 180 | 139 | 76 | 92 | HF |
| TSP10 | Female | 29 | 160 | 149 | 82.5 | 95.5 | Conventional |
| TSP11 | Male | 33 | 167 | 187 | 94.5 | 101.5 | Conventional |
| TSP12 | Male | 32 | 170 | 165 | 84.5 | 94 | HF |
| TSP13 | Male | 32 | 185 | 176 | 86.5 | 97 | Conventional |
| TSP14 | Female | 26 | 172 | 153 | 84 | 101.5 | HF |
| TSP15 | Female | 27 | 170 | 127 | 73 | 94 | HF |
| TSP16 | Female | 31 | 168 | 166 | 82 | 109 | HF |
| **Mean ± STD** | 8 Female | 26.3 ± 2.8 | 165.4 ± 4.9 | 135.3 ± 19.4 | 75.1 ± 6.6 | 93.8 ± 8.7 | |
| **Mean ± STD** | 8 Male | 32.8 ± 8.4 | 177.8 ± 8.8 | 169.3 ± 23.2 | 87.3 ± 8.7 | 95.6 ± 5.7 | |

We excluded individuals from this study if they were younger than 18 years of age, were pregnant, or had any of the following: implanted electronic devices, metallic implants in their torso and/or legs, a serious bone or blood disease or infection, heart disease such as arrhythmia, or a history of stroke, spinal cord injury or disease, or muscle or nerve impairments affecting the lower limbs. This study was approved by the Internal Review Board at Carnegie Mellon University (STUDY2020_00000452) and conducted in accordance with the Declaration of Helsinki. All participants signed a written informed consent form prior to their enrollment in the study. No participants had prior experience with tSCS.

### Study Protocol

#### EMG Electrode Placement

We prepared the skin of the left lower leg for recording using abrasive gel (Lemon Prep, Mavidon, USA), alcohol wipes (Braha Industries, USA) and conductive electrode gel (Signa Gel, Parker Laboratories BV, NL). All data were collected while the participant sat comfortably in a chair with both knees positioned at a 120° angle (Figure 1). We placed bipolar electromyography (EMG) electrodes (2 square 7/8”×7/8” Ag|AgCl foam electrodes; MVAP Medical Supplies, USA) approximately 1 cm apart on the soleus, lateral and medial gastrocnemius, and tibialis anterior muscles of the left leg. We secured a ground electrode (4×5 cm pregelled Ag|AgCl Natus electrode; MVAP Medical Supplies, USA) onto the left patella. EMG data were recorded using the SAGA64+ (TMSi, NL) at a sampling rate of 4000 Hz and streamed into MATLAB (MathWorks, USA) using custom software. Stimulation was delivered using a DS8R stimulator with a firmware update to allow frequencies up to 10 kHz (Digitimer, UK). The stimulator was triggered using a BNC 2090A connector accessory (National Instruments, USA) and custom MATLAB code.

**Figure 1.**
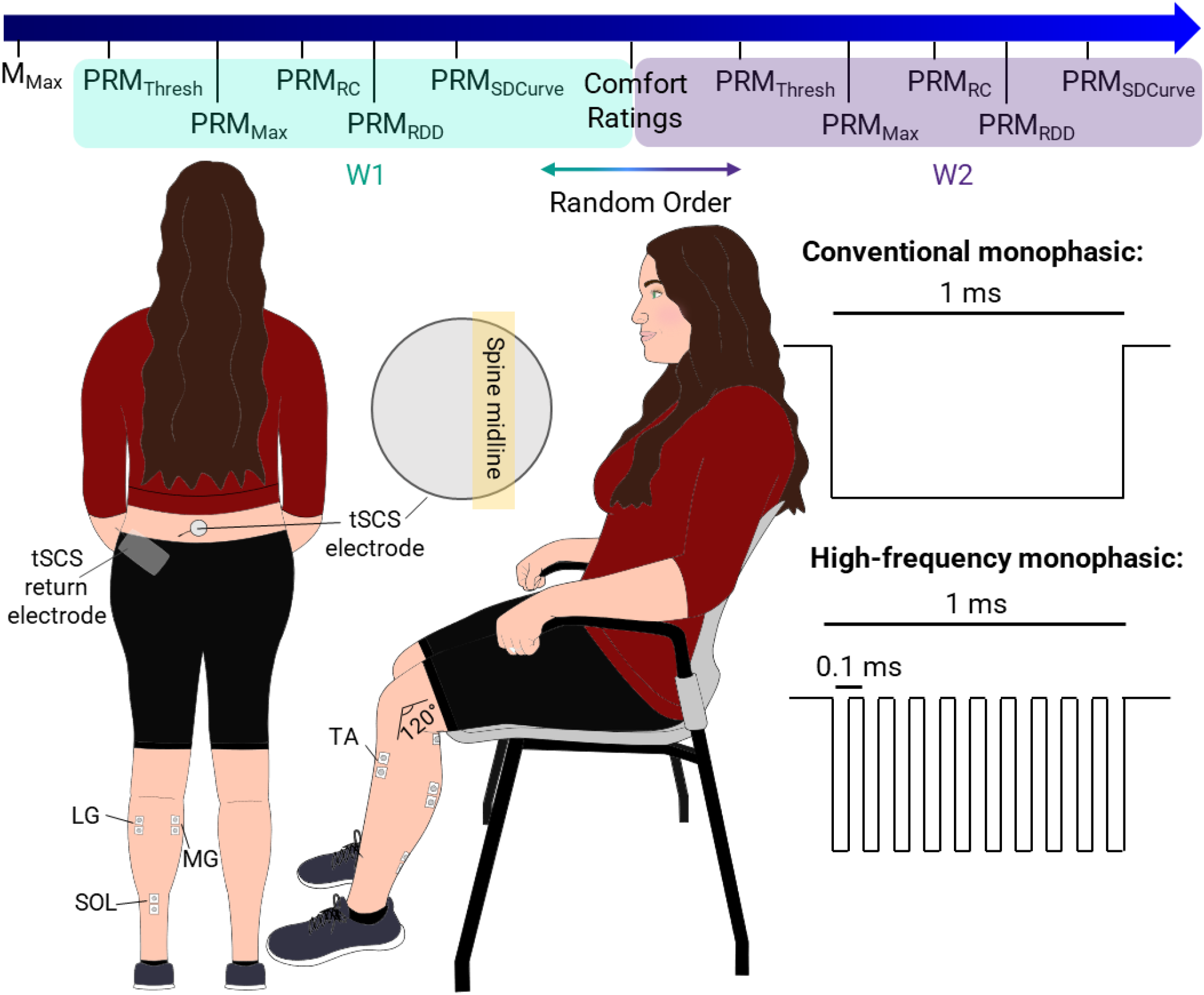
Experimental setup and procedures. The top bar shows the experimental timeline for collecting each measure. Waveform 1 (W1) and waveform 2 (W2) were either a conventional monophasic pulse or high-frequency burst pulse, the order of which was pseudo-randomized across participants. The bottom left shows the experimental setup with electrode placements. Electromyography (EMG) electrodes were placed on the soleus (SOL), lateral gastrocnemius (LG), medial gastrocnemius (MG), and tibialis anterior (TA) muscles. Transcutaneous spinal cord stimulation (tSCS) was delivered using a round electrode placed lateral to the T12-L1 spinal levels. The tSCS return electrode was placed on the anterior superior iliac spine. The inset of the round electrode shows the location of the electrode relative to the midline of the spine. The bottom right shows the two different waveforms studied. M_max_ = motor maximum; PRM = posterior root muscle; Thresh = threshold; Max = maximum; RC = recruitment curve; RDD = rate-dependent depression, SDCurve = strength-duration curve.

#### PRM Reflex Threshold

Participants were oblivious to the composition of the waveforms, and we pseudo randomized and balanced which one was delivered first.

We placed a round adhesive electrode for tSCS (3.2 cm diameter; ValuTrode, Axelgaard Manufacturing Co. Ltd., USA) paravertebrally left of the T12-L1 spinous processes, and a large rectangular electrode (7.5×13 cm, ValuTrode, Axelgaard Manufacturing Co. Ltd., USA) on the left anterior superior iliac spine. We wrapped the torso (6” Coban, 3M, USA) and placed a small piece of foam (12×17 cm) between the electrode and the back of the chair to maintain firm pressure on the stimulation site. Throughout the manuscript, we will refer to the monophasic pulse as the ‘conventional waveform’, and the pulse with the 10 kHz carrier frequency as the ‘high-frequency waveform’. Each of the following procedures were repeated for both waveforms.

We found the stimulation threshold for evoking a PRM reflex in the soleus muscle. We then determined the maximum PRM reflex amplitude (referred to as PRM_Max_), which was defined as the stimulation intensity at which the reflex amplitude no longer increased when stimulation intensity increased. In this study, we did not exceed a stimulation amplitude of 180 mA.

#### Recruitment Curves and Confirming Reflexes

We stimulated at 15 amplitudes between 5 mA below the PRM reflex threshold and 5-10 mA above the maximum intensity, in a random order. We delivered the stimuli 10 s apart and repeated each stimulation amplitude four times. To confirm that the evoked responses were reflexive, we tested for rate-dependent depression (RDD). We delivered a series of three pulses 10 s apart, and a series of three pulses 1 s apart, at a stimulation intensity approximately equal to 1.1-1.3 times PRM reflex threshold.

#### Varying Pulse Duration

The high-frequency waveform was active for half the time as the conventional waveform since the 10 kHz pulse train had a 50% duty cycle. Therefore, we sought to determine the effect of overall pulse duration on the PRM reflex threshold. We chose the following pulse durations: 100, 200, 300, 400, 500, 800, 1000, 1250, 1500, 1750, and 2000 µs. We determined PRM reflex threshold twice for each pulse duration and in a random order. It is important to note that for all other measures the pulse width was equal to 1 ms.

#### Discomfort Scoring

We asked the participants to rate their discomfort from the electrical stimulation at the electrode site on a visual analog scale from 0 to 10, where 0 indicated no discomfort at all, and 10 indicated the most uncomfortable sensation imaginable. This was done at each threshold value across the various pulse durations. Moreover, we asked participants to rank the strength of the contraction of the paraspinal muscles on a scale of 0 to 10, where 0 was no contraction at all, and 10 was the strongest contraction imaginable. Paraspinal contractions were rated across stimulation amplitudes delivered between 20 and 80 mA, at 15 mA intervals.

### Analysis and Statistics

We blanked the stimulus artefact in the EMG recordings by interpolating between points that occurred 4 ms prior to and 6 ms after the stimulus onset. We filtered EMG data using a second order band pass filter with cutoff frequencies of 20 Hz and 1999 Hz.

We determined the latency of the PRM reflexes as the time from the stimulation onset to the first inflection of the response. We constructed recruitment curves by calculating the average of the four PRM reflex responses elicited at each stimulation intensity. We then measured the peak-to-peak amplitude of the averaged responses according to stimulation charge. We found the slope of the recruitment curve by first using the MATLAB function *findchangepts* to find the inflection points. Then, we determined the slope of the line between those inflection points across the steepest part of the curve.

We created strength-duration curves for each waveform by finding the average threshold from the two repetitions at the same stimulation amplitude for each pulse duration. We created charge-duration curves by multiplying the threshold amplitude by the pulse duration, taking into account the duty cycle of the high-frequency waveform. We performed a linear regression on the charge-duration curves. From the linear regression, we extracted the slope, which is equal to the rheobase (Lin et al., 2002; Reilly et al., 1992) (also the threshold amplitude at infinite duration). The y-intercept is equivalent to the minimum charge threshold for short pulses (duration approaching zero). Finally, we calculated the chronaxie by taking the quotient of the y-intercept and the slope. The chronaxie is also equivalent to the duration for two times the rheobase from the strength-duration curve.

We obtained discomfort scores at threshold during as pulse duration was varied, and used the mean of the two scores for each pulse duration. There were no significant differences between the discomfort values across pulse durations (one-way analysis of variance (ANOVA); p = 1.0); therefore, we grouped the discomfort scores across pulse durations for comparisons.

We used paired t-tests to ascertain differences between the conventional and high-frequency waveforms for the following measures: stimulation amplitudes at PRM reflex threshold, peak-to-peak amplitudes at threshold, PRM reflex latencies, decrease in peak-to-peak amplitude following RDD, recruitment rate, and discomfort scores. We used the two one-sided t-test (Lakens, 2017; Rastogi, 2017) to test for equality between the two waveforms for the discomfort scores at threshold and latency of the PRM reflexes. We also tested for effects of order (starting with either the conventional or high-frequency waveform) on PRM reflex thresholds and comfort scores using paired t-tests. We performed a similar analysis with paired t-tests to determine if there were sex-based differences in PRM thresholds and comfort scores. Finally, we calculated the linear correlation coefficient between the threshold stimulation amplitude and the participant’s waist and hip circumferences.

## RESULTS

We evoked PRM reflexes in all lower-limb muscles that were tested in all participants using both waveforms. For the purpose of reporting our main findings, we focus on the soleus muscle only (Figure 2A, B) because the results from the other muscles were similar (Supplementary Figure 1). Near and above the PRM reflex threshold, large contractions in the paraspinal muscles occurred. The presence of paraspinal contractions was used as an indicator that the stimulation amplitude was likely to evoke a PRM reflex. Although we did not ask participants to describe paresthesias, over half (n = 10) of them reported experiencing paresthesias in their ankles to toes. Furthermore, we observed movements in both legs in half (n = 8) of the participants.

**Figure 2.**
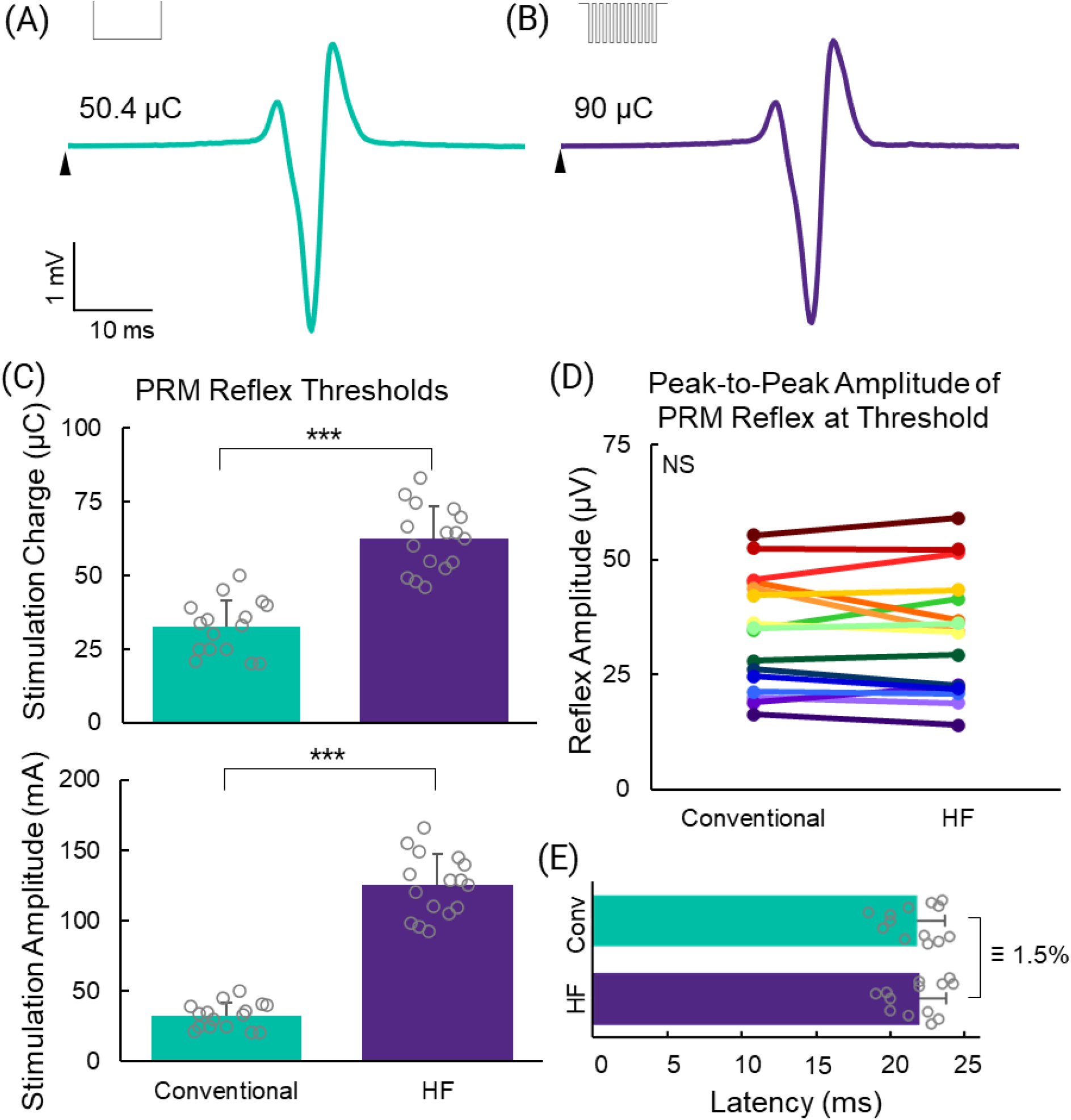
Posterior root muscle (PRM) reflexes from the soleus muscle of TSP08. Examples of PRM reflexes of the same peak-to-peak amplitude, evoked using the conventional (Conv) waveform at 50.4 µC (A) and the high-frequency (HF) waveform at 90µC (B). The waveforms are indicated by the insets. The onset of stimulation is indicated by an arrow. (C) Comparison of PRM reflex thresholds using the conventional versus the high-frequency waveform according to stimulation charge (top) and amplitude (bottom). (D) Comparison of the peak-to-peak amplitude of the PRM reflexes at threshold across all subjects. (E) Latency of the PRM reflexes. Error bars represent standard deviation; ***p < 0.001; NS = not significant; ≡1.5% indicates that the latencies were equivalent within a ±1.5% window.

### The High-Frequency Waveform Required More Charge to Evoke PRM Reflexes

The high-frequency waveform required approximately double the charge (62.5 ± 11.1 µC) to evoke a PRM reflex compared to the conventional waveform (32.4 ± 9.2 µC), which was significant (p < 0.001; Figure 2C). This equated to stimulation amplitudes of 125.1 ± 22.3 mA for the high-frequency waveform and 32.4 ± 9.2 mA for the conventional waveform. The peak-to-peak amplitude of the PRM reflexes evoked at threshold by the high-frequency waveform (33.1 ± 13.0) compared to the conventional waveform (34.8 ± 12.5) were not significantly different (p = 0.32; Figure 2D). The latencies of the PRM reflexes (21.8 ± 1.9 and 22.0 ± 1.8 for the conventional and high-frequency waveforms, respectively) were not significantly different (p = 0.11) and were equivalent with a window of ±1.5% (p = 0.034; Figure 2E).

Neither the waist nor hip circumferences correlated with PRM reflex thresholds for either waveform (R^2^ < 0.1; Supplementary Figure 2). Furthermore, sex did not influence the PRM reflex thresholds (p > 0.5; Supplementary Figure 4A). The order in which the waveforms were delivered also did not influence the PRM reflex threshold (p > 0.35; Supplementary Figure 4C).

### PRM Reflexes Exhibited Rate-Dependent Depression

We confirmed that the evoked responses were reflexive by observing RDD. When the stimuli were 10 s apart, the peak-to-peak amplitude of the PRM reflex remained constant (Figure 3A). When the inter-stimulus interval was 1 s, the peak-to-peak amplitude decreased with successive stimuli. This occurred for both waveforms (Figure 3B,C). On average, the peak-to-peak amplitude of the second PRM reflex evoked by the conventional and high-frequency waveforms were 47.5% (± 23.9%) and 64.9% (± 25.4%) of the first PRM reflex, respectively, and were not significantly different between waveforms (p = 0.09; Figure 3D). In most cases, the peak-to-peak amplitude of the third PRM reflex was also depressed; however, in 9/32 instances, the peak-to-peak amplitude of the third PRM reflex increased to approximately that of the first response (Supplementary Figure 3).

**Figure 3.**
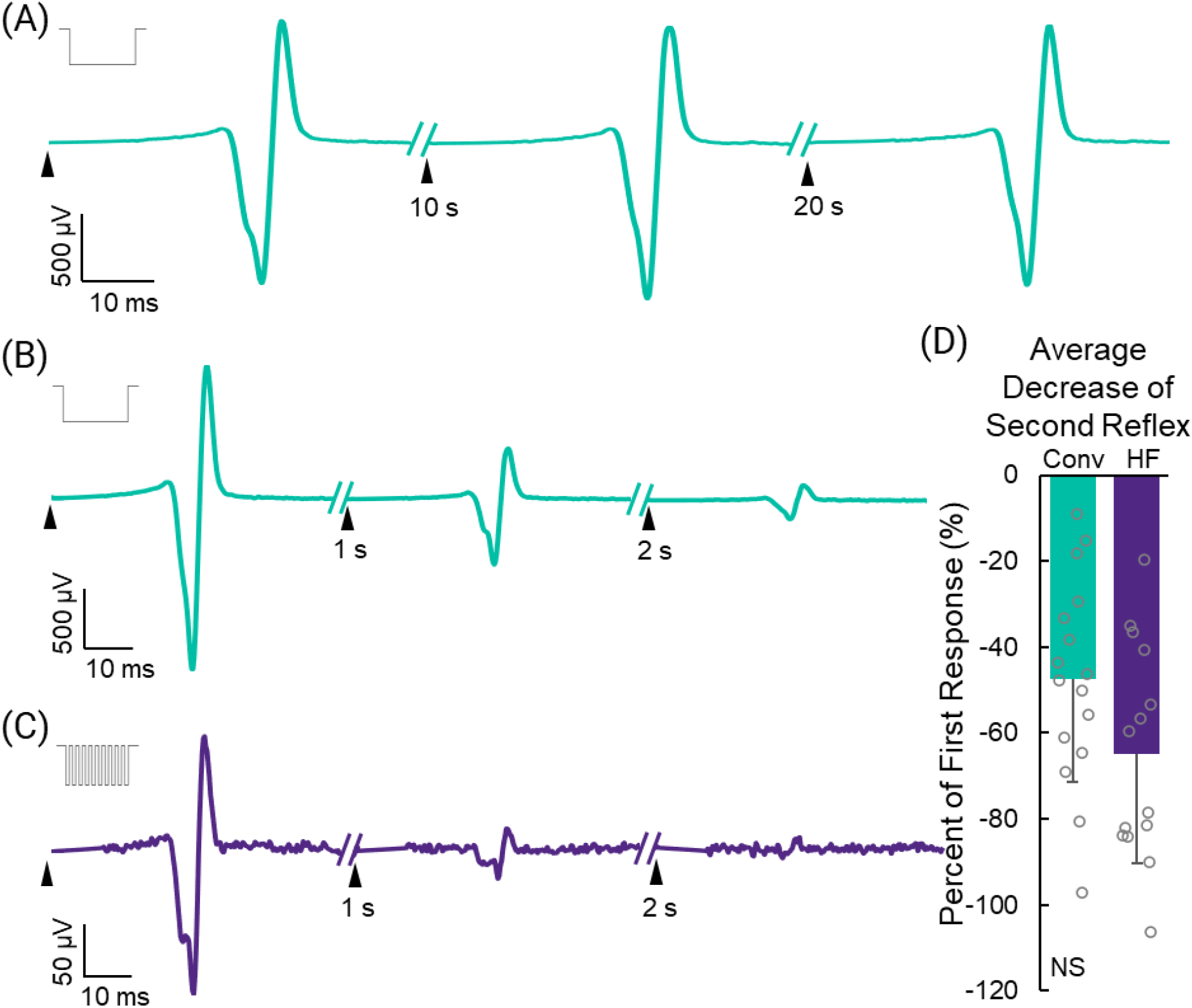
Rate-dependent depression. Example of posterior root muscle (PRM) reflexes following 3 consecutive stimulation pulses delivered 10 seconds apart (A), showing a constant peak-to-peak amplitude, or 1 second apart (B, C), showing a decrease in the peak-to-peak amplitude of the successive responses. Stimulation amplitudes were set to ∼1.1-1.3X PRM threshold. All responses shown were recorded in the soleus muscle from TSP02. The type of waveform used in each subfigure is indicated by the inset. The onset of stimulation is indicated by an arrow. (D) Average (- standard deviation) of the peak-to-peak amplitude of the second PRM reflex as a percentage of the first PRM reflex in the series. HF = high-frequency; Conv = conventional.

### Both Waveforms Had Similar Recruitment Rates

We collected recruitment curves by varying the stimulation amplitude between subthreshold and supramaximal amplitudes (Figure 4A,B,C). The slope of the recruitment curve indicates the rate of activation of the PRM reflex. In most participants, the PRM_Max_ was not reached using the high-frequency waveform. Therefore, the slope of the recruitment curve was calculated only if the high-frequency PRM_Max_ was at least 33% of the conventional PRM_Max_ (occurred only in TSP01, TSP02, TSP04, and TSP08). The recruitment rates for the conventional and high-frequency waveforms were 0.39 ± 0.29 mV/µC and 0.28 ± 0.23 mV/µC, respectively, and were not significantly different (p = 0.08).

**Figure 4.**
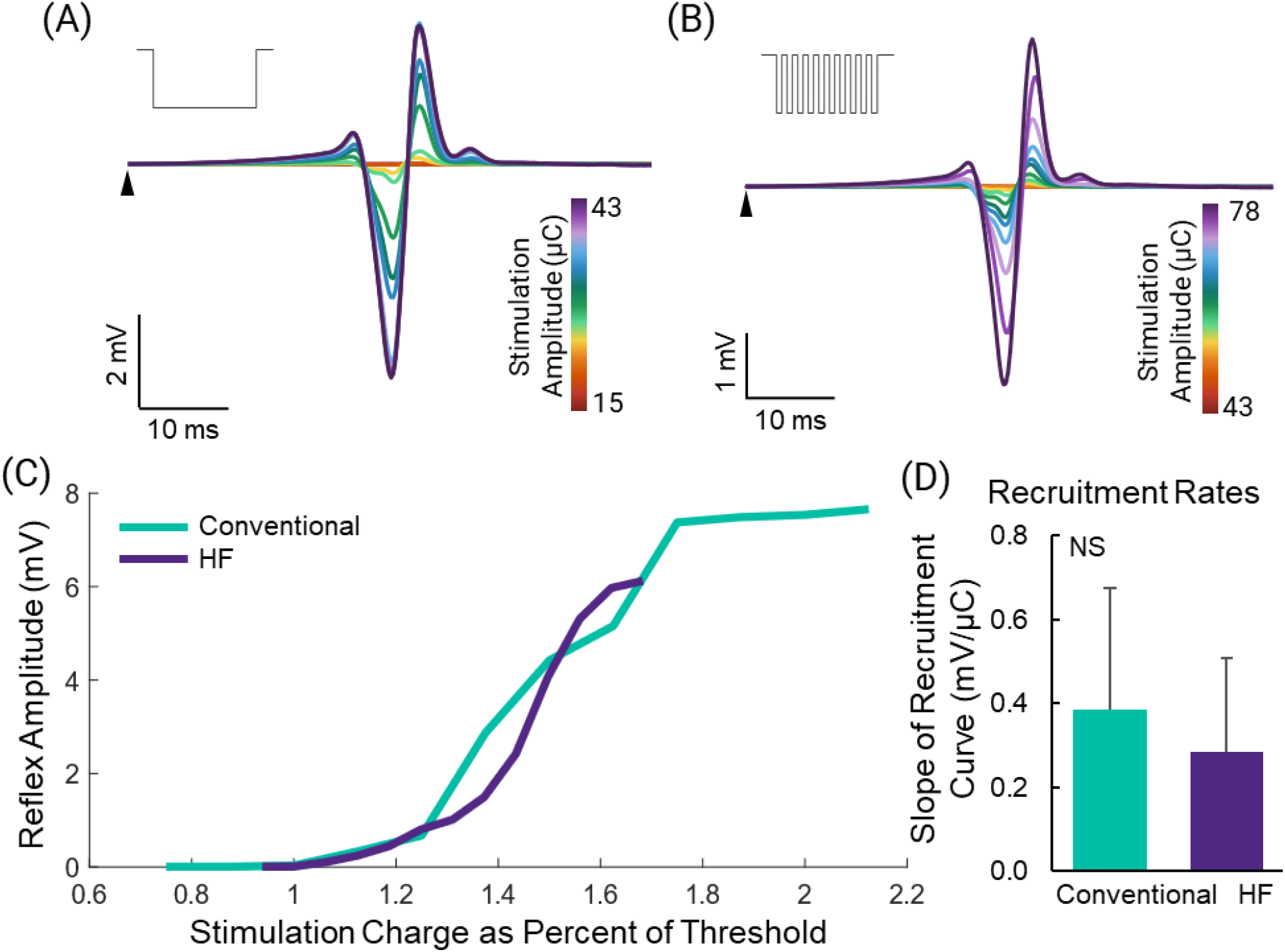
Recruitment of posterior root muscle (PRM) reflexes. PRM reflexes evoked across a range of amplitudes using the conventional waveform (A) and the high-frequency (HF) waveform (B). The type of waveform used in each subfigure is indicated by the inset. The onset of stimulation is indicated by an arrow. (C) Peak-to-peak amplitude of the PRM reflex response plotted against the stimulation charge as a percent of threshold. All responses shown were recorded in the soleus muscle from TSP02. (D) Recruitment rates of PRM reflexes for instances where the maximum peak-to-peak amplitude from the HF waveform was at least 33% of the maximum from the conventional waveform. Error bars represent standard deviation. ***p < 0.001; NS = not significant.

### The Waveforms Were Equally Comfortable at PRM Reflex Threshold

At PRM reflex threshold, the mean discomfort score for the conventional and high-frequency waveforms were 0.87 ± 0.2 and 1.03 ± 0.18, respectively, and were not significantly different from one another (p = 0.13), but were equivalent within a ±1% window (p = 0.002; Figure 5C). Sex did not have an effect on the discomfort scores (p > 0.25; Supplementary Figure 4B), nor did the order of which waveform was tested first (p > 0.55; Supplementary Figure 4D,E).

**Figure 5.**
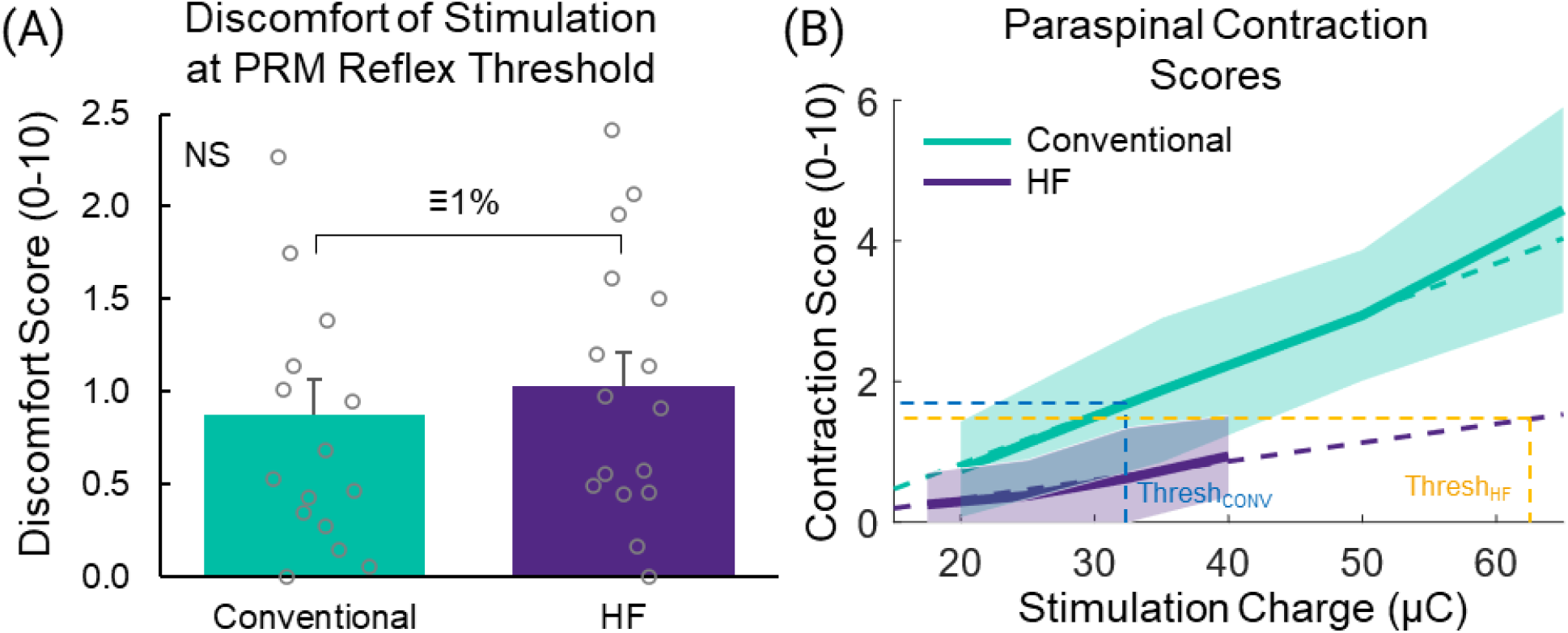
Discomfort and contraction scores. (A) Mean (+ standard deviation) of the scores for the discomfort felt at threshold along the strength-duration curves. (B) Mean (± standard deviation) scores for the paraspinal muscle contractions felt at each stimulation intensity for each waveform. NS = not significant; HF = high-frequency; CONV = conventional; ≡1% indicates that the discomfort scores were equivalent within a ±1% window.

**Figure 6.**
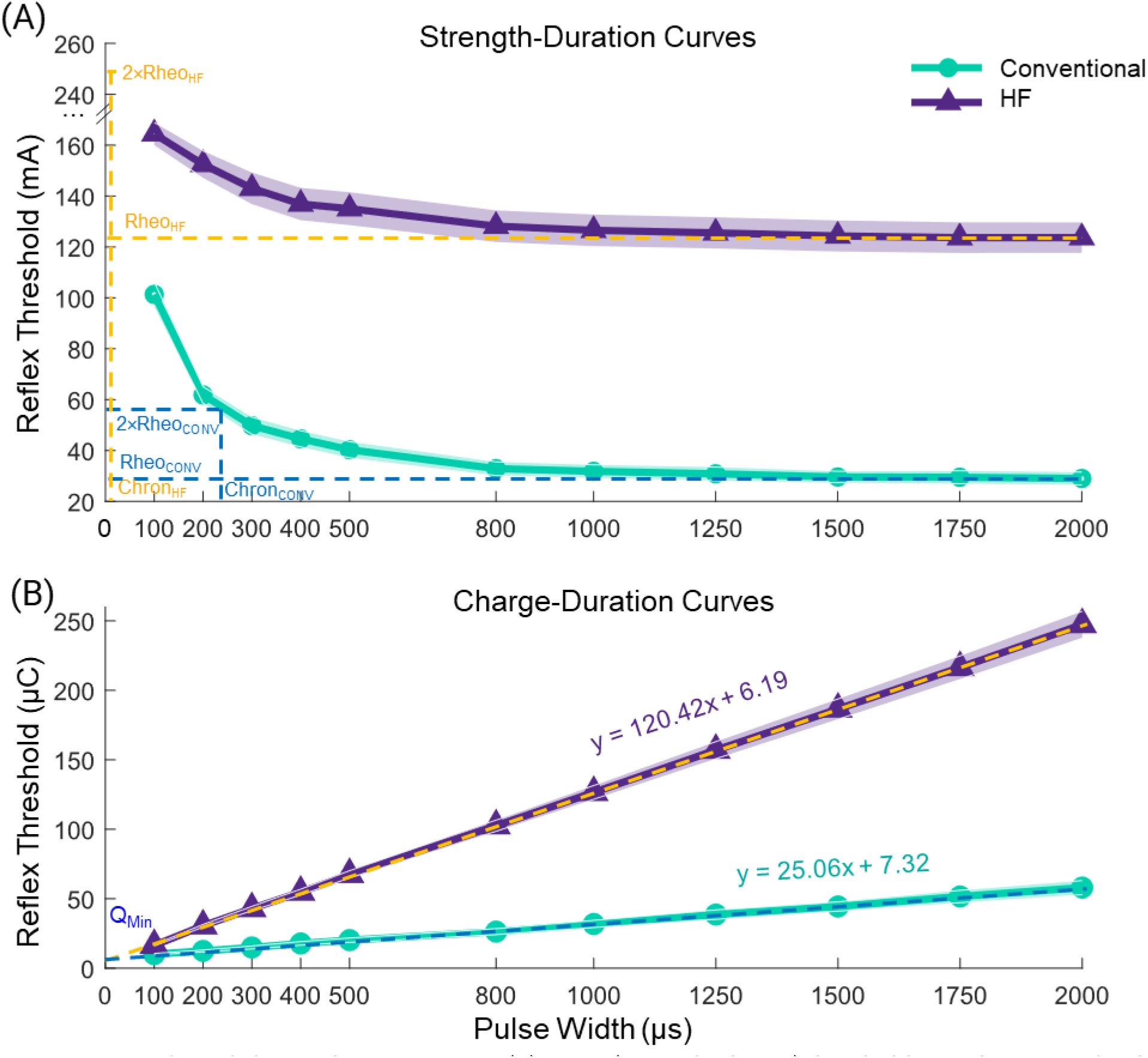
Strength- and charge-duration curves. (A) Mean (± standard error) threshold stimulation amplitude for evoking a posterior root muscle (PRM) reflex as the pulse duration was varied. Note: for the high-frequency (HF) waveform, the threshold for pulse durations < 300 µs were > 180 mA; therefore, the curve is incorrectly less steep. CONV = conventional waveform. (B) Mean (± standard error) charge threshold as pulse duration was varied. Q_Min_ = minimum charge; the equations are from linear regression.

As the stimulation charge increased, the contraction strength of paraspinal muscles were rated higher and increased linearly (conventional: R^2^ = 0.98; high-frequency: R^2^ = 0.96; Figure 5B). We used the equations from linear regression to estimate the contraction score at PRM reflex threshold:

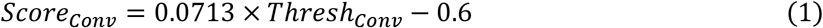

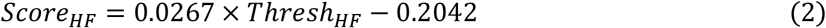

At threshold, the contraction score for the conventional and high-frequency waveforms were 1.71 and 1.46, respectively, indicating that they induced similarly strong paraspinal muscle contractions.

### The High-Frequency Waveform Delivered Charge Less Efficiently

The values for the rheobase, chronaxie, and minimum charge extracted from the charge-duration curves are listed in table 2 and labelled in figure 7. For 6 participants, the thresholds for narrower pulse durations were greater than 180 mA for the high-frequency waveform. In most of these cases, only the narrowest point (100 µs) was outside of this bound; however, in one instance, the 200 µs and 400 µs data points were outside this bound. Therefore, for pulse durations less than 400 µs, and in particular at 100 µs, the threshold value was underestimated, and these data points were removed from analysis. The strength-duration curve for the high-frequency waveform can be duplicated by shifting the strength-duration curve for the conventional waveform up by 94.7 mA (Supplementary Figure 5) in a near perfect overlap (Correlation = 96.1%). These results demonstrate that the high-frequency waveform activates the same afferent fibers as the conventional waveform, but less efficiently.

**Table 2.** Parameters from charge-duration and strength-duration curves. The parameters were determined using a linear regression of the charge-duration curves for each waveform.

| Parameter | Conventional Waveform | High-Frequency Waveform | HF/Conventional |
| --- | --- | --- | --- |
| Rheobase (mA) | 25.06 | 120.42 | 4.81 |
| Chronaxie (μs) | 292.1 | 51.4 | 0.18 |
| Minimum Charge (μC) | 7.32 | 6.19 | 0.85 |

## DISCUSSION

### PRM Reflexes

tSCS neuromodulation for the recovery of motor functions requires stimulation amplitudes at or near PRM reflex threshold to be effective (Hofstoetter et al., 2021; Inanici et al., 2021; Keller et al., 2021; Rath et al., 2018; Sayenko et al., 2019). PRM reflexes consist of a multi-segment superposition of H-reflexes and cutaneous afferent inputs (Freitas et al., 2022; Krenn et al., 2013; Minassian et al., 2007). A characteristic of spinal reflexes is the presence of RDD, indicated by the reduced amplitude of successive reflex responses following an initial pulse (Andrews et al., 2015). We observed RDD at 1 Hz, verifying that the responses were reflexive, similar to data from a previous study (Hofstoetter et al., 2019). We also reported periodic modulation of reflex amplitudes with further successive pulses. This behaviour has been reported previously with epidural SCS in people with SCI (Hofstoetter et al., 2015a). Our results suggest that both waveforms act through a similar reflex pathway because the shapes of the responses were identical within each participant and muscle, and they had similar recruitment characteristics.

### Discomfort of Stimulation at PRM Reflex Threshold

Discomfort is a limitation of electrical stimulation, especially at high amplitudes in people with intact sensation. Activation of cutaneous nociceptors and paraspinal muscle contractions are the main causes of discomfort during tSCS. Both types of discomfort have been reported in several studies, sometimes requiring discontinuation for several days (Hofstoetter et al., 2018; Keller et al., 2021; Manson et al., 2020). Our participants reported strong paraspinal muscle contractions and discomfort of electrical stimulation at higher amplitudes. However, therapeutic tSCS for motor recovery requires stimulation amplitudes at or below threshold (Hofstoetter et al., 2015b; Inanici et al., 2021; Rath et al., 2018), where mild paraspinal contractions are produced. Our results show that, at threshold, the strength of contractions of the paraspinal muscles are similar and small.

A recent study from the Sayenko lab investigated the maximum tolerable stimulation amplitude for tSCS with and without a 5 kHz carrier frequency (Manson et al., 2020). They noted that 70% of participants discontinued stimulation due to the discomfort of strong paraspinal contractions, 10% due to abdominal contractions (where their return electrode was located), and 20% due to stimulation-site discomfort. Participants could tolerate larger currents with high-frequency stimulation. However, when the maximum tolerable stimulation amplitude was normalized to the PRM reflex threshold, there was no difference between the waveforms. This showed that the reflex threshold and maximum tolerable stimulation amplitude scaled together with the addition of a high-frequency carrier. They concluded that high-frequency stimulation does not reduce discomfort at intensities needed to evoke PRM reflexes. Our results support this inference and explicitly demonstrate that, at PRM reflex threshold, the addition of a 10 kHz carrier frequency does not reduce discomfort.

### Mechanisms of High-Frequency Stimulation

A common claim to support the use of high-frequency tSCS is that it blocks local cutaneous afferents, resulting in painless stimulation (Gad et al., 2017; Y. Gerasimenko et al., 2015; Inanici et al., 2021; Manson et al., 2020; Sayenko et al., 2019). The concept of conduction block has been prevalent for decades. In the 1960’s, Tanner reported conduction block with 20 kHz stimulation, noting that large diameter fibers were blocked first (Tanner, 1962). A computational model showed that both deep and superficial large diameter afferents are activated with high-frequency stimulation (Medina and Grill, 2014). Lempka and colleagues used a computational model to show that high-frequency stimulation causes activation and blocking of axons, both in larger diameter fibers first (Lempka et al., 2015). Specifically, nerve fibers were activated at lower thresholds but blocked at amplitudes greater than the clinical range for SCS. It has also been demonstrated experimentally that 10 kHz stimulation blocked large diameter afferents at lower amplitudes than for unmyelinated fibers, which required supraclinical amplitudes (Joseph and Butera, 2011). Blocking may dominate when high-frequency stimulation is delivered tonically (Bhadra et al., 2018; Ward et al., 2004), but only after eliciting a large initial response (Ackermann et al., 2011). tSCS neuromodulation delivers stimulation in bursts of high-frequency trains, typically at 30-50 Hz, which may have different blocking properties than tonic stimulation. If blocking does occur during tSCS, the large diameter fibers would be blocked before small diameter fibers. Given that single bursts of high-frequency stimulation are equally as comfortable as a conventional pulse and the known blocking properties of the different afferent fibers, the notion that high-frequency tSCS blocks local cutaneous afferents but activates deep spinal roots is likely incorrect.

Over half our participants experienced paresthesias in their lower limbs using either waveform. Paresthesias are thought to indicate activation of Aβ mechanoreceptors (Caylor et al., 2019; Rogers et al., 2022) and have been reported in previous tSCS studies using the conventional waveform (Hofstoetter et al., 2021, 2015b). High-frequency stimulation is used in some forms of SCS for pain treatment. In this embodiment, high-frequency stimulation is referred to as ‘paresthesia-free’ (Caylor et al., 2019; Sdrulla et al., 2018); however, SCS is applied tonically, not in bursts of trains like tSCS. Whether or not Aβ fibers are activated using paresthesia-free SCS remains unclear (Crosby et al., 2017; Freitas et al., 2022; Rogers et al., 2022; Sdrulla et al., 2018; Song et al., 2014)(Freitas et al., 2022; Rogers et al., 2022). The presence of paresthesias and the question of blocking indicate that further studies are needed to differentiate the fibers being activated not only by different waveforms, but also when they are applied in single pulses, trains, or continuously.

Strength-duration curves describe activation of excitable tissues, where less excitable tissues are found higher and to the right of more excitable tissues. Here, the high-frequency strength-duration curve was a direct copy of the conventional strength-duration curve, shifted up but not right, suggesting that both waveforms activated the same afferent fibers. The rheobase indicates the absolute minimum current required to activate a nerve fiber, and the chronaxie is a function of the axon membrane time constant, which corresponds to excitability (Lin et al., 2002; Reilly et al., 1992). The rheobase for the high-frequency waveform was nearly 5 times greater than the conventional waveform, and the chronaxie more than 5 times narrower. High-frequency stimulation causes nerve excitation through summation, where each brief pulse in the burst brings the membrane closer to threshold, eventually leading to an action potential (Ward et al., 2004; Ward and Lucas-Toumbourou, 2007). Since the high-frequency waveform had a 50% duty cycle, one would expect the rheobase to be twice that of the conventional waveform. Our results show that integration does occur over the duration of the high-frequency pulse, but the summation is leaky, requiring significantly higher levels of current to achieve threshold than is required of the conventional waveform. This leaky summation results in less efficient excitation of afferent fibers and excessive current consumption (Irnich, 1980).

### Study Limitations

The upper limit for the stimulation current in this study was 180 mA. We chose this limit based on other tSCS studies. A few papers tested stimulation amplitudes up to 200 mA in people with SCI (Gad et al., 2018, 2017; Y. Gerasimenko et al., 2015); a recent study stimulated as high as 1000 mA in neurologically intact individuals without adverse events (Manson et al., 2020). The primary focus of our study was to investigate the comfort of each waveform at threshold for evoked PRM reflexes; therefore, the behaviour at higher stimulation amplitudes was less critical.

### Implications for Future Studies and Clinical Translation

Therapeutic SCS and tSCS use trains of stimulation pulses, often applied at 30-50 Hz, which is quite different than the single pulse applied in this study. A recent study reported that continuous trains of tSCS (30 Hz) were tolerated at only 15% of the current for a single pulse, which was 56% of motor threshold (Manson et al., 2020). This seems to contradict the numerous reports of using tSCS neuromodulation in people with intact sensation at just below (Hofstoetter et al., 2021, 2020; Inanici et al., 2021; Rath et al., 2018) or above (Gad et al., 2018; Y. Gerasimenko et al., 2015) motor threshold. We as a field need to understand the recruitment properties and comfort of single-pulse and continuous trains of tSCS to better understand how it engages with spinal reflex pathways during neuromodulation. Similar studies should be undertaken for other neuromodulation methods as well.

## CONCLUSIONS

tSCS neuromodulation for motor recovery requires stimulation amplitudes near PRM reflex threshold to engage spinal reflex pathways. High-frequency stimulation is less efficient and requires more charge to evoke PRM reflexes than a conventional monophasic stimulation pulse. At reflex thresholds, high-frequency stimulation is not more comfortable than conventional stimulation, despite several claims of pain-free stimulation when a high-frequency carrier is used. While tSCS offers a non-invasive approach to neuromodulation of motor output in the spinal cord, results from this study discourage using a high-frequency carrier because it leads to unnecessarily high levels of charge to be applied.

## ACKNOWLEDGMENTS

Thank you to Axelgaard for providing us with a suite of electrode samples to complete this work. We would also like to thank our research subjects for their commitment to science and willingness to participate in this study. We appreciate the feedback we received from our Data Safety and Monitoring Board throughout the completion of this study.

## AUTHOR CONTRIBUTIONS

AND, MC, and DJW conceived of the study; AND and DJW designed the study; AND collected and analyzed all data; CAH and MGK analyzed portions of the data; AND, MC, and DJW interpreted the data; AND, CAH, and MGK created the figures; AND wrote the first draft of the manuscript; all authors refined and approved the final manuscript prior to submission.

## SUPPLEMENTARY FIGURES

**Supplementary Figure 1.**
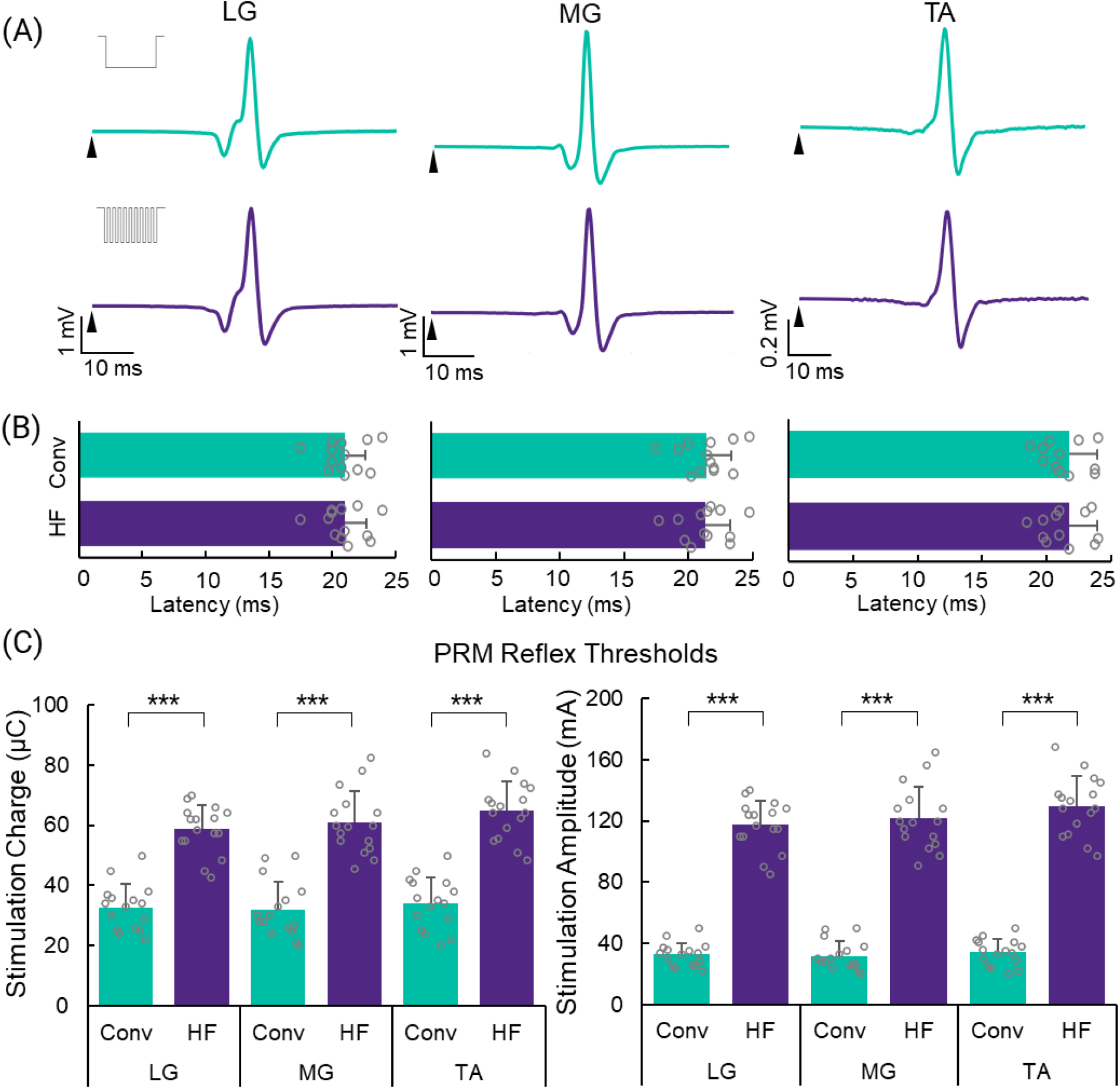
Posterior root-muscle (PRM) reflexes in other lower leg muscles. (A) Examples of identical PRM reflexes evoked by each waveform (indicated by inset) for the lateral gastrocnemius (LG), medial gastrocnemius (MG), and tibialis anterior (TA) muscles. The onset of stimulation is indicated by an arrow. (B) Latency of PRM reflexes for each muscle and waveform. (C) PRM reflex thresholds for each muscle and waveform expressed in stimulation charge (left) and current (right). HF = high-frequency; Conv = conventional; error bars represent standard deviation; ***p < 0.001.

**Supplementary Figure 2.**
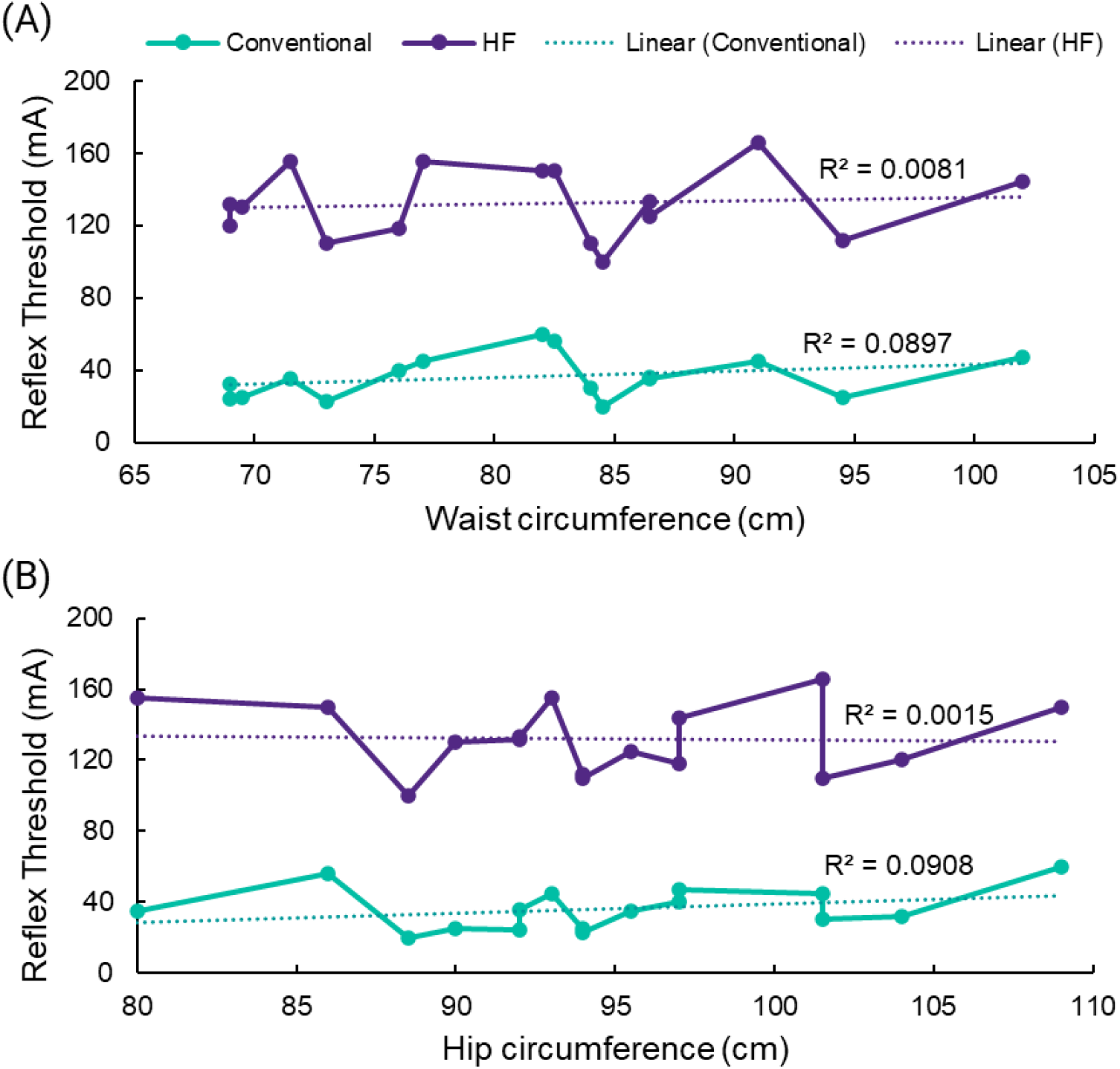
Effect of size on thresholds. Posterior root-muscle (PRM) reflex threshold as a function of waist circumference (A) and hip circumference (B). Linear correlations are shown by the dotted lines.

**Supplementary Figure 3.**
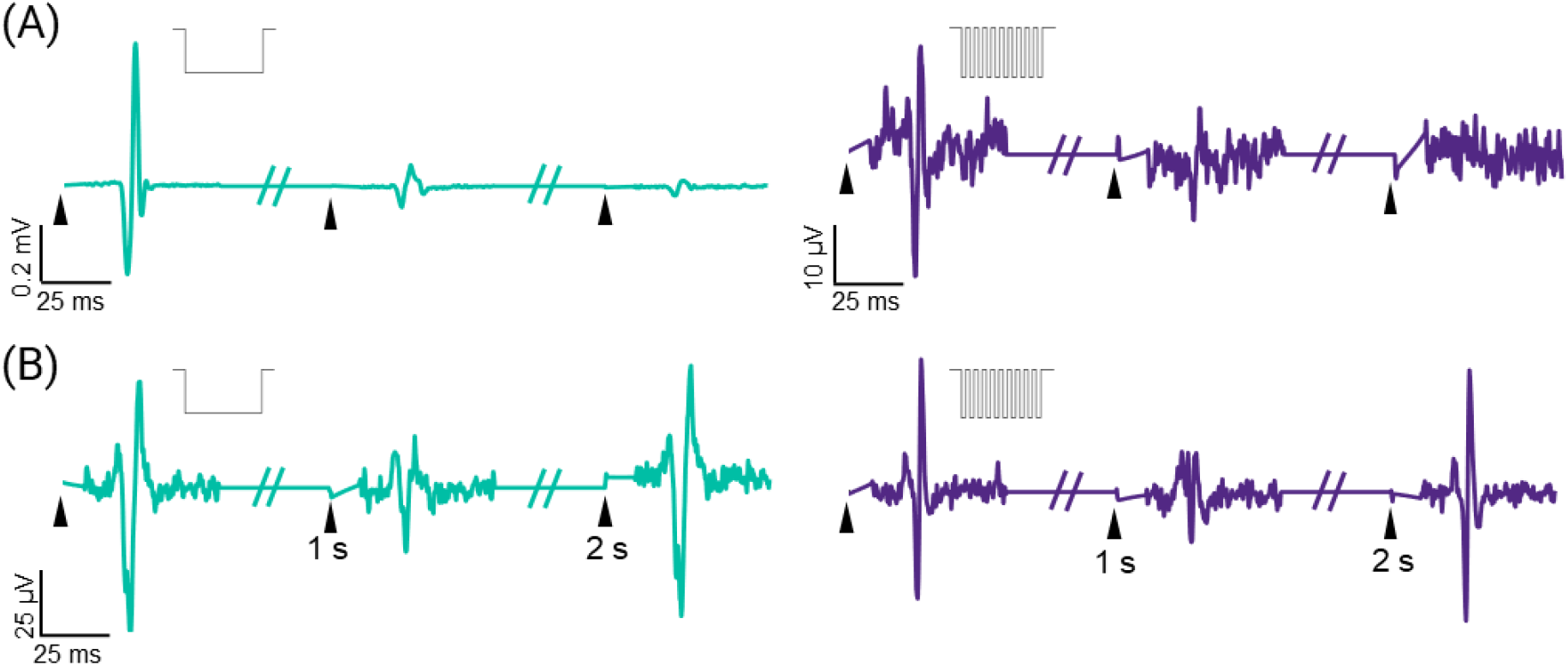
Additional examples of rate-dependent depression of posterior root-muscle (PRM) reflexes. (A) Successive depression of PRM reflexes in the lateral gastrocnemius muscle in TSP02. (B) Periodic modulation of PRM reflexes in the soleus muscles of TSP16 (left) and TSP12 (right). The type of waveform used in each subfigure is indicated by the inset. The onset of stimulation is indicated by an arrow.

**Supplementary Figure 4.**
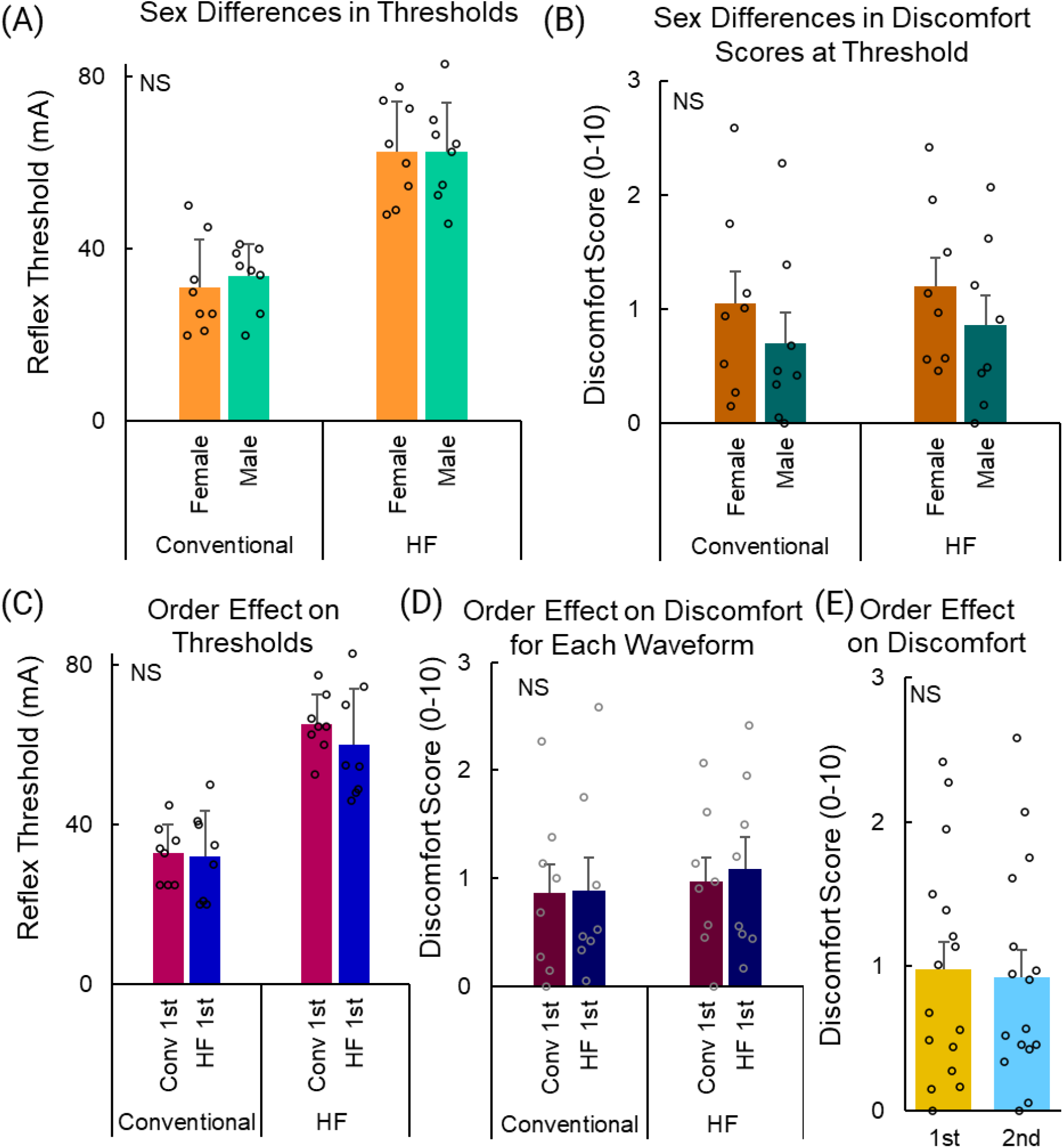
Effects of sex and order on thresholds and comfort. Effect of sex on posterior root-muscle (PRM) reflex threshold (A) and discomfort scores (B). Effect of order (conventional waveform or high-frequency (HF) waveform first) on PRM reflex threshold (C) and discomfort scores (D). (E) Effect of order (first versus second waveform) on discomfort scores. NS = not significant; error bars represent standard deviation.

**Supplementary Figure 5.**
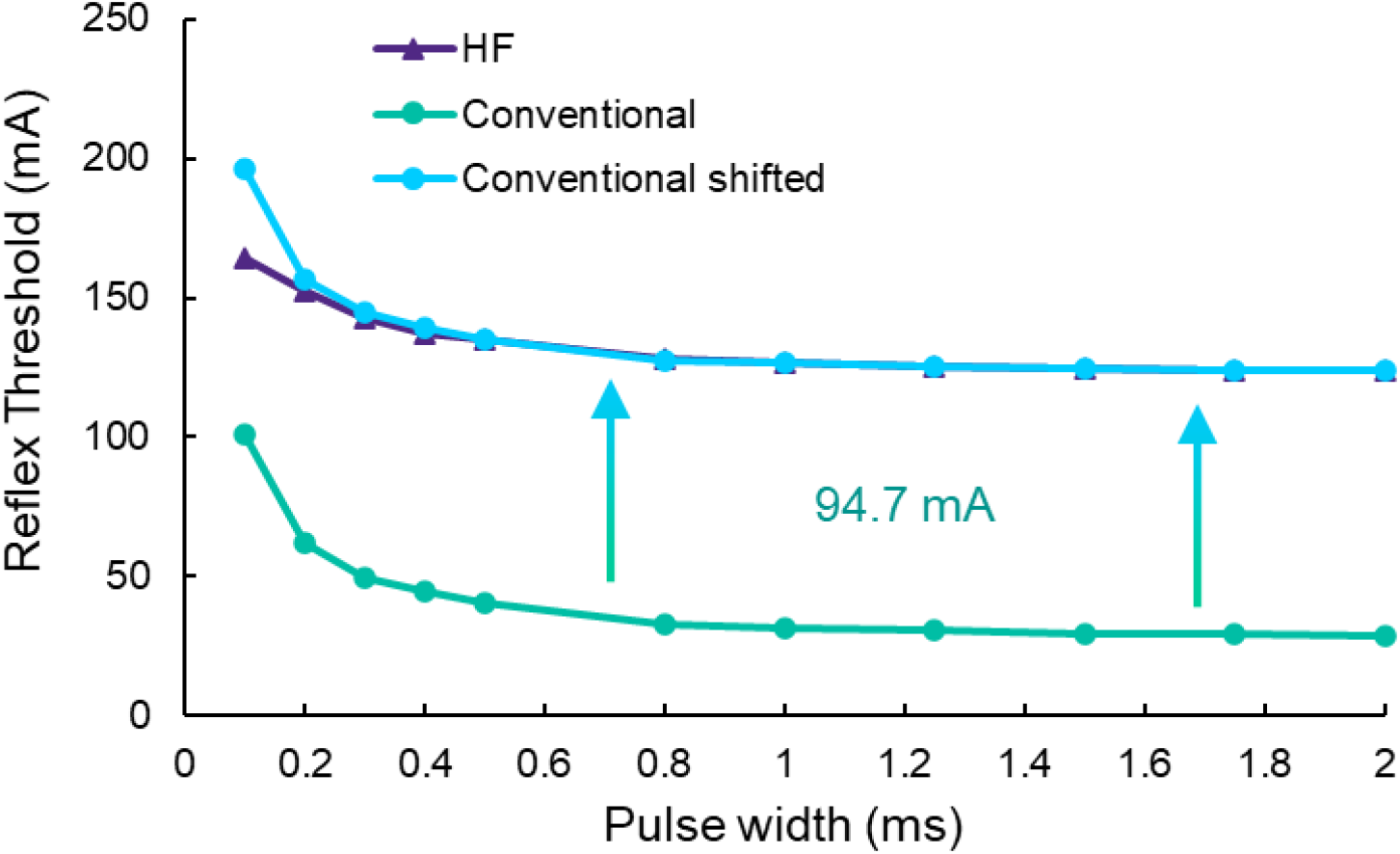
Strength-duration curve shift. We shifted the strength-duration curve of the conventional waveform up by 94.7 mA to overlap with the strength-duration curve of the high-frequency (HF) waveform to demonstrate their likeness. Note: for the HF waveform, the threshold for pulse durations < 300 µs were > 180 mA; therefore, the curve is incorrectly less steep.

## REFERENCES

Ackermann DM, Bhadra N, Foldes EL, Kilgore KL. 2011. Conduction block of whole nerve without onset firing using combined high frequency and direct current. Med Biol Eng Comput 49:241–251. doi:10.1007/s11517-010-0679-x

Andrews JC, Stein RB, Roy FD. 2015. Post-activation depression in the human soleus muscle using peripheral nerve and transcutaneous spinal stimulation. Neurosci Lett 589:144–149. doi:10.1016/j.neulet.2015.01.041

Angeli CA, Boakye M, Morton RA, Vogt J, Benton K, Chen Y, Ferreira CK, Harkema SJ. 2018. Recovery of Over-Ground Walking after Chronic Motor Complete Spinal Cord Injury. N Engl J Med 379:1244– 1250. doi:10.1056/NEJMoa1803588

Bhadra Narendra, Foldes E, Vrabec T, Kilgore K, Bhadra Niloy. 2018. Temporary persistence of conduction block after prolonged kilohertz frequency alternating current on rat sciatic nerve. J Neural Eng 15:016012. doi:10.1088/1741-2552/aa89a4

Capogrosso M, Wenger N, Raspopovic S, Musienko P, Beauparlant J, Bassi Luciani L, Courtine G, Micera S. 2013. A computational model for epidural electrical stimulation of spinal sensorimotor circuits. J Neurosci Off J Soc Neurosci 33:19326–19340. doi:10.1523/JNEUROSCI.1688-13.2013

Carhart MR, He J, Herman R, D’Luzansky S, Willis WT. 2004. Epidural spinal-cord stimulation facilitates recovery of functional walking following incomplete spinal-cord injury. IEEE Trans Neural Syst Rehabil Eng Publ IEEE Eng Med Biol Soc 12:32–42. doi:10.1109/TNSRE.2003.822763

Caylor J, Reddy R, Yin S, Cui C, Huang M, Huang C, Ramesh R, Baker DG, Simmons A, Souza D, Narouze S, Vallejo R, Lerman I. 2019. Spinal cord stimulation in chronic pain: evidence and theory for mechanisms of action. Bioelectron Med 5. doi:10.1186/s42234-019-0023-1

Crosby ND, Janik JJ, Grill WM. 2017. Modulation of activity and conduction in single dorsal column axons by kilohertz-frequency spinal cord stimulation. J Neurophysiol 117:136–147. doi:10.1152/jn.00701.2016

Freitas RM de, Capogrosso M, Nomura T, Milosevic M. 2022. Preferential activation of proprioceptive and cutaneous sensory fibers compared to motor fibers during cervical transcutaneous spinal cord stimulation: a computational study. J Neural Eng 19:036012. doi:10.1088/1741-2552/ac6a7c

Freyvert Y, Yong NA, Morikawa E, Zdunowski S, Sarino ME, Gerasimenko Y, Edgerton VR, Lu DC. 2018. Engaging cervical spinal circuitry with non-invasive spinal stimulation and buspirone to restore hand function in chronic motor complete patients. Sci Rep 8:1–10. doi:10.1038/s41598-018-33123-5

Gad P, Gerasimenko Y, Edgerton VR. 2019. Tetraplegia to Overground Stepping Using Non-Invasive Spinal Neuromodulation2019 9th International IEEE/EMBS Conference on Neural Engineering (NER). Presented at the 2019 9th International IEEE/EMBS Conference on Neural Engineering (NER). pp. 89–92. doi:10.1109/NER.2019.8717096

Gad P, Gerasimenko Y, Zdunowski S, Turner A, Sayenko D, Lu DC, Edgerton VR. 2017. Weight Bearing Over-ground Stepping in an Exoskeleton with Non-invasive Spinal Cord Neuromodulation after Motor Complete Paraplegia. Front Neurosci 11. doi:10.3389/fnins.2017.00333

Gad P, Lee S, Terrafranca N, Zhong H, Turner A, Gerasimenko Y, Edgerton VR. 2018. Non-Invasive Activation of Cervical Spinal Networks after Severe Paralysis. J Neurotrauma 35:2145–2158. doi:10.1089/neu.2017.5461

Gerasimenko Y, Gad P, Sayenko D, McKinney Z, Gorodnichev R, Puhov A, Moshonkina T, Savochin A, Selionov V, Shigueva T, Tomilovskaya E, Kozlovskaya I, Edgerton VR. 2016. Integration of sensory, spinal, and volitional descending inputs in regulation of human locomotion. J Neurophysiol 116:98– 105. doi:10.1152/jn.00146.2016

Gerasimenko Y, Gorodnichev R, Moshonkina T, Sayenko D, Gad P, Reggie Edgerton V. 2015. Transcutaneous electrical spinal-cord stimulation in humans. Ann Phys Rehabil Med 58:225–231. doi:10.1016/j.rehab.2015.05.003

Gerasimenko YP, Lu DC, Modaber M, Zdunowski S, Gad P, Sayenko DG, Morikawa E, Haakana P, Ferguson AR, Roy RR, Edgerton VR. 2015. Noninvasive Reactivation of Motor Descending Control after Paralysis. J Neurotrauma 32:1968–1980. doi:10.1089/neu.2015.4008

Gill ML, Grahn PJ, Calvert JS, Linde MB, Lavrov IA, Strommen JA, Beck LA, Sayenko DG, Van Straaten MG, Drubach DI, Veith DD, Thoreson AR, Lopez C, Gerasimenko YP, Edgerton VR, Lee KH, Zhao KD. 2018. Neuromodulation of lumbosacral spinal networks enables independent stepping after complete paraplegia. Nat Med. doi:10.1038/s41591-018-0175-7

Harkema S, Gerasimenko Y, Hodes J, Burdick J, Angeli C, Chen Y, Ferreira C, Willhite A, Rejc E, Grossman RG, Edgerton VR. 2011. Effect of epidural stimulation of the lumbosacral spinal cord on voluntary movement, standing, and assisted stepping after motor complete paraplegia: a case study. Lancet 377:1938–1947. doi:10.1016/S0140-6736(11)60547-3

Hofstoetter US, Danner SM, Freundl B, Binder H, Mayr W, Rattay F, Minassian K. 2015a. Periodic modulation of repetitively elicited monosynaptic reflexes of the human lumbosacral spinal cord. J Neurophysiol 114:400–410. doi:10.1152/jn.00136.2015

Hofstoetter US, Freundl B, Binder H, Minassian K. 2019. Recovery cycles of posterior root-muscle reflexes evoked by transcutaneous spinal cord stimulation and of the H reflex in individuals with intact and injured spinal cord. PloS One 14:e0227057. doi:10.1371/journal.pone.0227057

Hofstoetter US, Freundl B, Binder H, Minassian K. 2018. Common neural structures activated by epidural and transcutaneous lumbar spinal cord stimulation: Elicitation of posterior root-muscle reflexes. PloS One 13:e0192013. doi:10.1371/journal.pone.0192013

Hofstoetter US, Freundl B, Danner SM, Krenn MJ, Mayr W, Binder H, Minassian K. 2020. Transcutaneous Spinal Cord Stimulation Induces Temporary Attenuation of Spasticity in Individuals with Spinal Cord Injury. J Neurotrauma 37:481–493. doi:10.1089/neu.2019.6588

Hofstoetter US, Freundl B, Lackner P, Binder H. 2021. Transcutaneous Spinal Cord Stimulation Enhances Walking Performance and Reduces Spasticity in Individuals with Multiple Sclerosis. Brain Sci 11:472. doi:10.3390/brainsci11040472

Hofstoetter US, Hofer C, Kern H, Danner SM, Mayr W, Dimitrijevic MR, Minassian K. 2013. Effects of transcutaneous spinal cord stimulation on voluntary locomotor activity in an incomplete spinal cord injured individual. Biomed Eng Biomed Tech. doi:10.1515/bmt-2013-4014

Hofstoetter US, Krenn M, Danner SM, Hofer C, Kern H, McKay WB, Mayr W, Minassian K. 2015b. Augmentation of Voluntary Locomotor Activity by Transcutaneous Spinal Cord Stimulation in Motor-Incomplete Spinal Cord-Injured Individuals. Artif Organs 39:E176–186. doi:10.1111/aor.12615

Hofstoetter US, McKay WB, Tansey KE, Mayr W, Kern H, Minassian K. 2014. Modification of spasticity by transcutaneous spinal cord stimulation in individuals with incomplete spinal cord injury. J Spinal Cord Med 37:202–211. doi:10.1179/2045772313Y.0000000149

Inanici F, Brighton LN, Samejima S, Hofstetter CP, Moritz CT. 2021. Transcutaneous Spinal Cord Stimulation Restores Hand and Arm Function After Spinal Cord Injury. IEEE Trans Neural Syst Rehabil Eng 29:310–319. doi:10.1109/TNSRE.2021.3049133

Inanici F, Samejima S, Gad P, Edgerton VR, Hofstetter CP, Moritz CT. 2018. Transcutaneous Electrical Spinal Stimulation Promotes Long-Term Recovery of Upper Extremity Function in Chronic Tetraplegia. IEEE Trans Neural Syst Rehabil Eng Publ IEEE Eng Med Biol Soc 26:1272–1278. doi:10.1109/TNSRE.2018.2834339

Irnich W. 1980. The chronaxie time and its practical importance. Pacing Clin Electrophysiol PACE 3:292– 301. doi:10.1111/j.1540-8159.1980.tb05236.x

Joseph L, Butera RJ. 2011. High-frequency stimulation selectively blocks different types of fibers in frog sciatic nerve. IEEE Trans Neural Syst Rehabil Eng Publ IEEE Eng Med Biol Soc 19:550–557. doi:10.1109/TNSRE.2011.2163082

Keller A, Singh G, Sommerfeld JH, King M, Parikh P, Ugiliweneza B, D’Amico J, Gerasimenko Y, Behrman AL. 2021. Noninvasive spinal stimulation safely enables upright posture in children with spinal cord injury. Nat Commun 12:5850. doi:10.1038/s41467-021-26026-z

Knikou M, Murray LM. 2019. Repeated transspinal stimulation decreases soleus H-reflex excitability and restores spinal inhibition in human spinal cord injury. PloS One 14:e0223135. doi:10.1371/journal.pone.0223135

Krenn M, Toth A, Danner SM, Hofstoetter US, Minassian K, Mayr W. 2013. Selectivity of transcutaneous stimulation of lumbar posterior roots at different spinal levels in humans. Biomed Tech (Berl) 58 Suppl 1. doi:10.1515/bmt-2013-4010

Lakens D. 2017. Equivalence Tests: A Practical Primer for t Tests, Correlations, and Meta-Analyses. Soc Psychol Personal Sci 8:355–362. doi:10.1177/1948550617697177

Lempka SF, McIntyre CC, Kilgore KL, Machado AG. 2015. Computational analysis of kilohertz frequency spinal cord stimulation for chronic pain management. Anesthesiology 122:1362–1376. doi:10.1097/ALN.0000000000000649

Lin CS-Y, Chan JHL, Pierrot-Deseilligny E, Burke D. 2002. Excitability of human muscle afferents studied using threshold tracking of the H reflex. J Physiol 545:661–669. doi:10.1113/jphysiol.2002.026526

Manson GA, Calvert JS, Ling J, Tychhon B, Ali A, Sayenko DG. 2020. The relationship between maximum tolerance and motor activation during transcutaneous spinal stimulation is unaffected by the carrier frequency or vibration. Physiol Rep 8:e14397. doi:10.14814/phy2.14397

Medina LE, Grill WM. 2014. Volume conductor model of transcutaneous electrical stimulation with kilohertz signals. J Neural Eng 11:066012. doi:10.1088/1741-2560/11/6/066012

Minassian K, Persy I, Rattay F, Dimitrijevic MR, Hofer C, Kern H. 2007. Posterior root-muscle reflexes elicited by transcutaneous stimulation of the human lumbosacral cord. Muscle Nerve 35:327–336. doi:10.1002/mus.20700

Powell MP, Verma N, Sorensen E, Carranza E, Boos A, Fields D, Roy S, Ensel S, Barra B, Balzer J, Goldsmith J, Friedlander RM, Wittenberg G, Fisher LE, Krakauer JW, Gerszten PC, Pirondini E, Weber DJ, Capogrosso M. 2022. Epidural stimulation of the cervical spinal cord improves voluntary motor control in post-stroke upper limb paresis. doi:10.1101/2022.04.11.22273635

Rastogi A. 2017. Two One-Sided Test (TOST) for equivalence.

Rath M, Vette AH, Ramasubramaniam S, Li K, Burdick J, Edgerton VR, Gerasimenko YP, Sayenko DG. 2018. Trunk Stability Enabled by Noninvasive Spinal Electrical Stimulation after Spinal Cord Injury. J Neurotrauma 35:2540–2553. doi:10.1089/neu.2017.5584

Reilly JP, Antoni H, Chilbert MA, Skuggevig W, Sweeney JD. 1992. Electrical Stimulation and Electropathology. Cambridge University Press.

Rogers ER, Zander HJ, Lempka SF. 2022. Neural Recruitment During Conventional, Burst, and 10-kHz Spinal Cord Stimulation for Pain. J Pain 23:434–449. doi:10.1016/j.jpain.2021.09.005

Rowald A, Komi S, Demesmaeker R, Baaklini E, Hernandez-Charpak SD, Paoles E, Montanaro H, Cassara A, Becce F, Lloyd B, Newton T, Ravier J, Kinany N, D’Ercole M, Paley A, Hankov N, Varescon C, McCracken L, Vat M, Caban M, Watrin A, Jacquet C, Bole-Feysot L, Harte C, Lorach H, Galvez A, Tschopp M, Herrmann N, Wacker M, Geernaert L, Fodor I, Radevich V, Van Den Keybus K, Eberle G, Pralong E, Roulet M, Ledoux J-B, Fornari E, Mandija S, Mattera L, Martuzzi R, Nazarian B, Benkler S, Callegari S, Greiner N, Fuhrer B, Froeling M, Buse N, Denison T, Buschman R, Wende C, Ganty D, Bakker J, Delattre V, Lambert H, Minassian K, van den Berg CAT, Kavounoudias A, Micera S, Van De Ville D, Barraud Q, Kurt E, Kuster N, Neufeld E, Capogrosso M, Asboth L, Wagner FB, Bloch J, Courtine G. 2022. Activity-dependent spinal cord neuromodulation rapidly restores trunk and leg motor functions after complete paralysis. Nat Med 28:260–271. doi:10.1038/s41591-021-01663-5

Samejima S, Caskey CD, Inanici F, Shrivastav SR, Brighton LN, Pradarelli J, Martinez V, Steele KM, Saigal R, Moritz CT. 2022. Multisite Transcutaneous Spinal Stimulation for Walking and Autonomic Recovery in Motor-Incomplete Tetraplegia: A Single-Subject Design. Phys Ther 102:pzab228. doi:10.1093/ptj/pzab228

Sayenko DG, Rath M, Ferguson AR, Burdick JW, Havton LA, Edgerton VR, Gerasimenko YP. 2019. Self-Assisted Standing Enabled by Non-Invasive Spinal Stimulation after Spinal Cord Injury. J Neurotrauma 36:1435–1450. doi:10.1089/neu.2018.5956

Sdrulla AD, Guan Y, Raja SN. 2018. Spinal Cord Stimulation: Clinical Efficacy and Potential Mechanisms. Pain Pract 18:1048–1067. doi:10.1111/papr.12692

Shealy CN, Mortimer JT, Reswick JB. 1967. Electrical inhibition of pain by stimulation of the dorsal columns: preliminary clinical report. Anesth Analg 46:489–491.

Song Z, Viisanen H, Meyerson BA, Pertovaara A, Linderoth B. 2014. Efficacy of Kilohertz-Frequency and Conventional Spinal Cord Stimulation in Rat Models of Different Pain Conditions. Neuromodulation Technol Neural Interface 17:226–235. doi:10.1111/ner.12161

Tanner JA. 1962. Reversible blocking of nerve conduction by alternating-current excitation. Nature 195:712–713. doi:10.1038/195712b0

Ward AR, Lucas-Toumbourou S. 2007. Lowering of sensory, motor, and pain-tolerance thresholds with burst duration using kilohertz-frequency alternating current electric stimulation. Arch Phys Med Rehabil 88:1036–1041. doi:10.1016/j.apmr.2007.04.009

Ward AR, Robertson VJ. 1998. Sensory, motor, and pain thresholds for stimulation with medium frequency alternating current. Arch Phys Med Rehabil 79:273–278. doi:10.1016/s0003-9993(98)90006-5

Ward AR, Robertson VJ, Ioannou H. 2004. The effect of duty cycle and frequency on muscle torque production using kilohertz frequency range alternating current. Med Eng Phys 26:569–579. doi:10.1016/j.medengphy.2004.04.007

Ward AR, Shkuratova N. 2002. Russian Electrical Stimulation: The Early Experiments. Phys Ther 82:1019– 1030. doi:10.1093/ptj/82.10.1019

